# Integrating regulatory information via combinatorial control of the transcription cycle

**DOI:** 10.1101/039339

**Authors:** Clarissa Scholes, Angela H. DePace, Álvaro Sánchez

## Abstract

Combinatorial regulation of gene expression by multiple transcription factors (TFs) enables cells to carry out sophisticated computations that are key to cellular decision-making. How is the information contained in multiple TF binding sites integrated to dictate the rate of transcription? The dominant model is that direct or indirect physical interactions between TFs enable them to combinatorially recruit each other and RNA polymerase to the promoter. Here we develop a quantitative framework to explore an alternative model, where combinatorial gene regulation can result from TFs working on different kinetic steps of the transcription cycle. Our results clarify the null hypotheses for independent action of TFs and show that combinatorial control of the transcription cycle can generate a wide range of analog and Boolean computations without requiring the input regulators to be simultaneously co-localized in the nucleus. This work emphasizes the importance of deciphering TF function beyond activation and repression, highlights the role of the basal promoter in processing regulatory information and suggests qualitative explanations for the flexibility of regulatory evolution.

## INTRODUCTION

To control gene expression, multiple transcription factors (TFs) bind to clusters of their cognate binding sites in regulatory DNA; this relays information from environmental and developmental cues into gene expression outcomes. Combinatorial regulation enables a small number of TFs to implement different types of computations (Struhl 1991; Carey 1998) in different cell types or over developmental time (Spitz & Furlong 2012). A grand challenge is to decipher how information contained in multiple binding sites—or more broadly, in multiple regulatory elements in a gene locus—is integrated to quantitatively determine the rate of transcription (Figure 1A,B).

**Figure 1.**
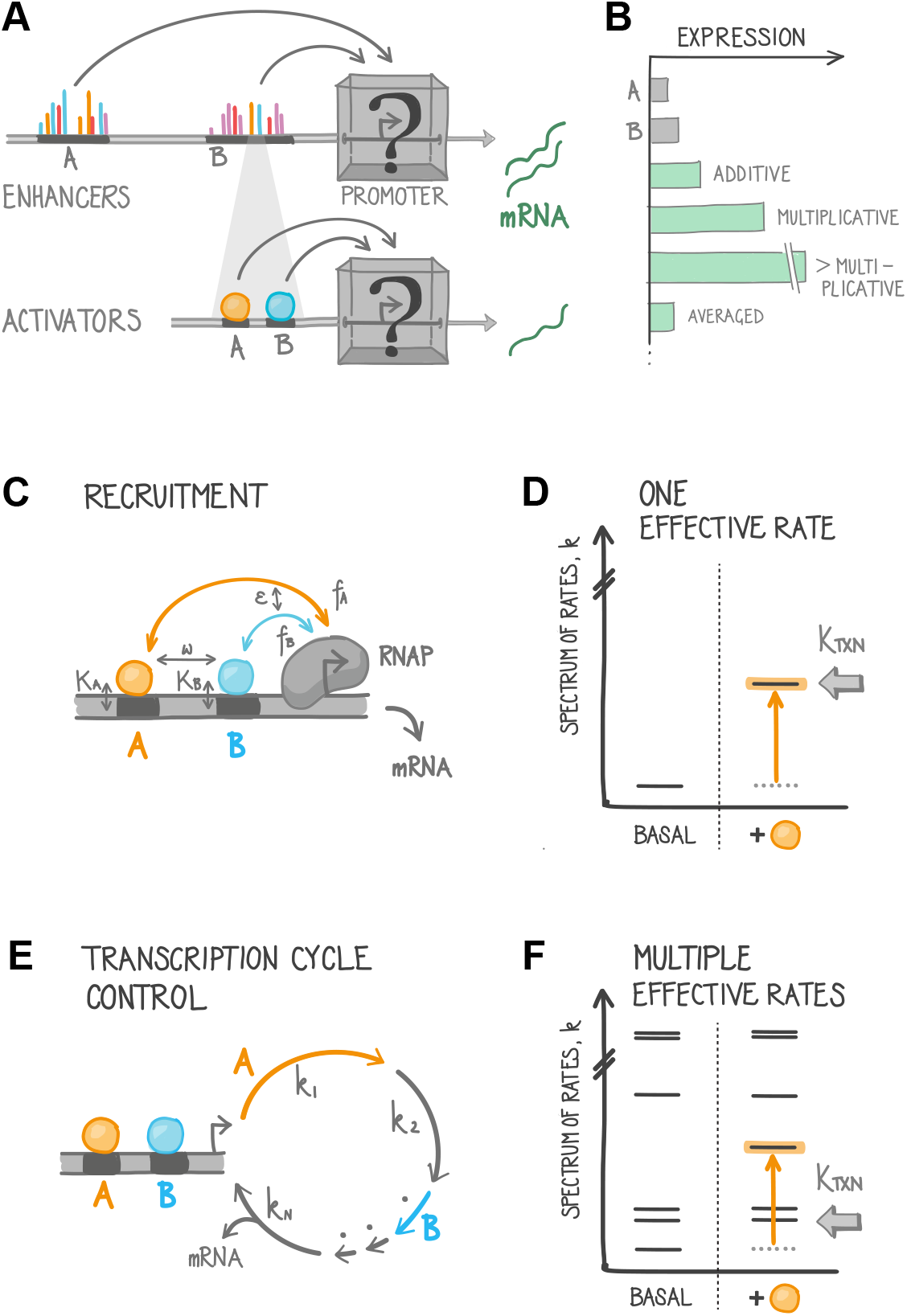
Promoters integrate information from multiple regulatory sequences. **(A)** Multiple regulatory sequences (top) or individual bound transcriptional activators (bottom) impinge on a given promoter at the same time. Together these regulatory sequences determine the level and timing of gene expression. We ask how these inputs are integrated by the promoter, by defining the overall expression level driven by two elements (either enhancers or activators) as a function of their individual effects. **(B)** Two regulatory elements, A and B, may combine in different ways to determine the overall expression level. **(C)** Models of combinatorial transcriptional control usually assume that the single limiting step is “recruitment” of RNA Polymerase (RNAP) to the promoter. Recruitment is mediated by physical interactions between different transcription factors (TFs, A and B) and the DNA, and between TFs and RNAP. The rate of transcription is assumed to be proportional to RNAP occupancy of the promoter. *K_A_* – dissociation constant for TF A. *f_A_* – strength of interaction between TF A and RNAP. *ω* – cooperativity between TFs A and B in binding to DNA. *ε* - cooperativity between the TFs in recruiting RNAP. **(D)** Recruitment models implicitly assume a single slow effective rate in transcription. Transcription is limited by the dissociation of RNAP from the promoter, and activators work by reducing this dissociation rate. The single effective rate is described by the rate constant *k* (first column). Adding an activator (orange) accelerates this effective rate, raising the overall rate of transcription (*k_TXN_*, arrow). All other steps in transcription are assumed to be infinitely fast (not shown). **(E)** The transcription cycle contains numerous distinct steps (Box 1). Shown here are the effective rates of *N* arbitrary steps, with shorter arrows indicating faster rates. TFs may work on the same or different steps in the cycle. Here, TF A accelerates effective rate *k_1_* (orange) while B accelerates one of the faster, unlabeled rates (blue). **(F)** The effective rates of the transcription cycle are depicted before and after addition of an activator (orange) that works on the slowest step. The overall rate of transcription increases when the activator is added, but is limited by the next slowest rate.

To analyze how computations are carried out by TFs binding to regulatory DNA, computational models of combinatorial gene regulation have largely focused on how TFs interact, cooperatively or competitively, to recruit or repel one another and the basal transcriptional machinery (Bintu et al. 2005a; He et al. 2010; Veitia 2003; Coulon et al. 2013; Wang et al. 1999; Frank et al. 2012; Gertz et al. 2009). A key implicit assumption of these “recruitment” models (Ptashne & Gann 1997) is that there is a single effective rate-limiting step in transcription (Figure 1D). It follows from this assumption that all TFs act by modulating this rate-limiting step. At many bacterial genes, where RNA polymerase (RNAP) is in direct competition with a repressor whose binding site overlaps its own, the approximation of a single rate limiting step may be reasonable (Garcia & Phillips 2011; Brewster et al. 2012). It should be noted, however, that this is just an approximation, and that many bacterial promoters do not conform to it (Box 1). A frequent additional assumption is that TF and RNAP binding can be calculated at equilibrium, thus this class of models is often referred to as “thermodynamic” (Bintu et al. 2005a). Several recent reviews give practical guidelines on building and applying these models (Bintu et al. 2005a; Bintu et al. 2005b; Sherman & Cohen 2012; Dresch et al. 2013).

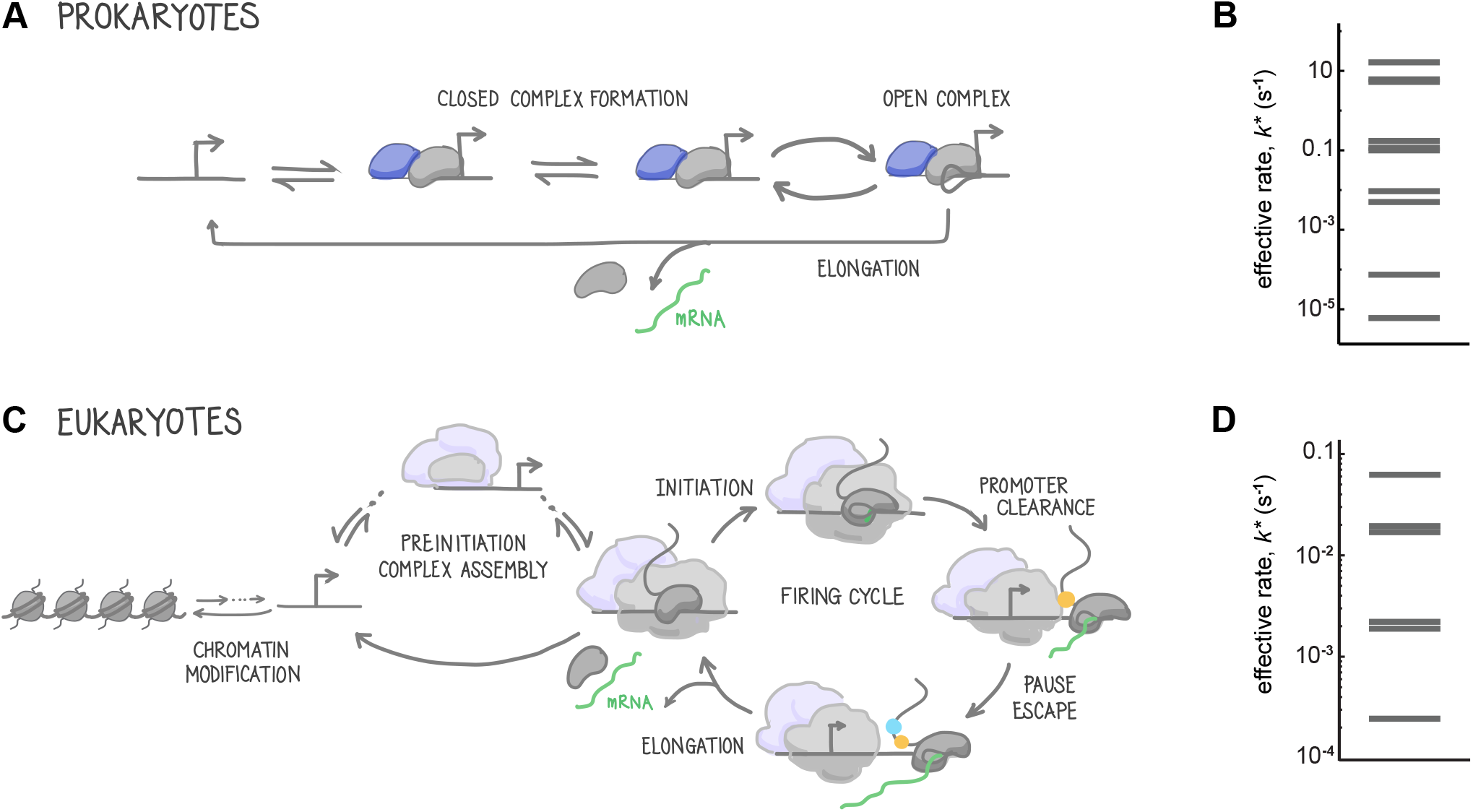
Box 1. The transcription cycle in prokaryotes and eukaryotes. (A) Prokaryotes. RNAP is recruited to the promoter through association with a sigma factor to form the closed complex (Gross et al. 1998; Browning & Busby 2004). Subsequent melting of the DNA forms the open complex (McClure 1985) and RNAP then escapes the promoter into elongation. It was widely assumed that recruitment was the rate-limiting step in bacterial transcription (Ptashne & Gann 1997); however, the transition from initiation to elongation is rate limiting at many promoters (Reppas et al. 2006; Wade & Struhl 2008). Diagram of a bacterial transcription cycle adapted from Friedman & Gelles (2012). (B) We used the forward and reverse reaction rates measured in Friedman & Gelles (2012) to calculate the “effective” forward rates through the transcription cycle (Results and Figure S1A-C). As this rates diagram shows, assuming a single slow step in transcription may be problematic even for bacterial promoters: the two slowest effective rates are within an order of magnitude of one another. (C) Eukaryotes. The transcription cycle begins with the formation of the preinitiation complex comprised of RNAP and general TFs (Fuda et al. 2009). The chromatin surrounding the promoter is modified, which is coordinated by DNA-bound TFs that recruit chromatin remodelers and nucleosome modifying enzymes. Transcription initiation can take place once the DNA has been unwound to form the open complex. RNAP must subsequently escape the grip of proteins at the promoter to enter into elongation. At most genes in higher eukaryotes, RNAP pauses downstream of the promoter and must be actively released from this paused state into productive elongation through the body of the gene (reviewed in (Zeitlinger et al. 2007; Jonkers et al. 2014; Adelman & Lis 2012)). Diagram of higher eukaryotic transcription cycle adapted from Fuda et al. (2009) and Boettiger 2013. (D) The “effective” rates for a simplified eukaryotic transcription cycle were calculated from Stasevich et al (2014) (Results and Figure S1D-F).

The assumption that there is only a single rate-limiting step in transcription may not, however, be appropriate for eukaryotes. The eukaryotic transcription cycle is composed of numerous distinct steps, from clearing of nucleosomes around the promoter to release of paused RNAP (Box 1), and various of these steps can be rate-limiting (Fuda et al. 2009; Green 2005; Nechaev & Adelman 2011; Jonkers et al. 2014; Hahn 2004; Choubey et al. 2015). Analysis of transcriptional dynamics in eukaryotic cells indicate that there are multiple slow biochemical reactions required to turn a gene on, and that at the promoters studied their rates must be within the same order of magnitude (Coulon et al. 2013; Zoller et al. 2015; Choubey et al. 2015).

If two or more steps have similar rates, control of transcription might not be dominated by regulation of a single step (Figure 1F); this could allow for combinatorial control by regulators working on different steps, as proposed over 20 years ago by Herschlag & Johnson (1993). Indeed, TFs are known to work on distinct steps in the transcription cycle (Fuda et al. 2009; Blau et al. 1996; Govind et al. 2005; Stasevich et al. 2014; Yankulov et al. 1994; Brown et al. 1998; Liu et al. 2003). For example, the activator SP1 works primarily on initiation (Blau et al. 1996) and Tat targets elongation (Barboric & Peterlin 2005); many (perhaps most) TFs affect multiple steps (e.g. Gcn4 (Govind et al. 2005) and CRP (Liu et al. 2003)). Since it was proposed, some evidence consistent with kinetic control of the transcription cycle has come from a small number of experimental studies (Blau et al. 1996; Gonzalez-Couto et al. 1997; Keung et al. 2014; Blair et al. 1996). For example, Bentley and colleagues saw greater-than-additive expression only between pairs of activation domains that worked on different steps (Blau et al. 1996).

Here, we aim to provide a synthesis of these theoretical and experimental observations by exploring combinatorial control via regulation of different slow steps in the transcription cycle. We emphasize that this type of kinetic control is compatible with combinatorial control arising from cooperative interactions between TFs in recruiting RNAP; they are not mutually exclusive mechanisms. For the sake of clarity, here we focus on a quantitative model of kinetic control as this mechanism is relatively under explored. Quantitative models enable us to reason about complex processes and to make falsifiable predictions that can be tested experimentally (Gunawardena 2014; Phillips 2015). For transcription, such models are also essential for drawing mechanistic interpretations from increasing volumes of high-throughput quantitative data that attempt to empirically characterize regulatory DNA (e.g. Sharon et al. 2012; Smith et al. 2013; Patwardhan et al. 2012). Our goal is here is to probe the regulatory outcomes and computations that emerge when cells combinatorially regulate different kinetic steps. We show that kinetic control of the transcription cycle can be used to implement a wide range of logical and analog computations, without requiring any cooperative interactions between input TFs. The pairwise interactions between TFs that are needed to produce many of these computations under recruitment-based models require that the TFs are simultaneously co-localized in the nucleus; combinatorial control of the transcription cycle can recapitulate these computations without the requirement for TF co-localization.

## RESULTS

### Calculating transcription from a cycle with no internal loops

To give intuition for how regulation of a transcription cycle is affected by all of its component steps, we first derive a closed form expression for a cycle with no internal loops. Here, the rate of transcription is the inverse of the sum of the transition times between states, but when reversible steps are included, the *effective time* taken to complete each step depends on the reverse as well as the forward transition times. We summarise the effect of a reverse rate between *state j* and *state j-1* by introducing a parameter that captures the average number of times the promoter switches backwards; we call this the number of ‘Rejections’ from *state j,*

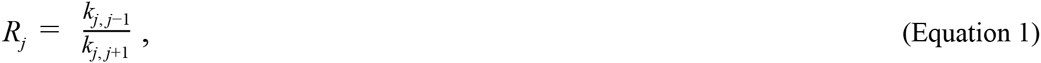

where *k_j, j+1_* is the rate of the forward reaction from *state j* to the subsequent state, *j + 1* and *k_j, j−1_* is the rate of the backward reaction from *state j* to the previous state, *j-1* The effective time taken to complete step *j* (i.e. the “handling time” of that state) depends on the number of Rejections from all of the *downstream* reversible transitions in the transcription cycle. For example, for a cycle comprised of *N =* 4 states in which the first three are connected by reversible transitions and the final step in which mRNA is produced is irreversible, we calculate the effective transition times, denoted by *T_j_\** for *j = 2:N* as follows, where τ*_j_* is the time taken to transition between states *j* and *j+1*:

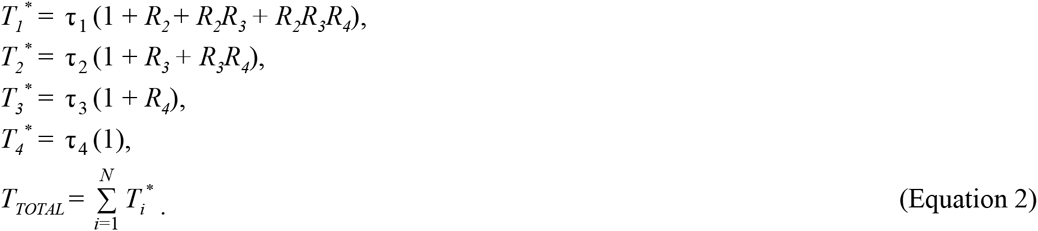

The total time taken to complete a cycle with reversible and irreversible steps (Equation 2) can be written in compact form for a cycle with an arbitrary number of steps, *N*, as:

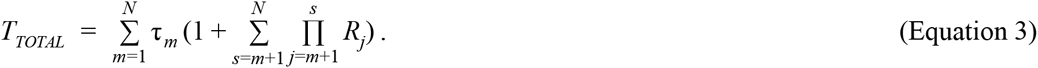

The rate of transcription is the inverse of *T_TOTAL_*. For the previous example of a four-step cycle, what we end up with is actually ten terms (all the terms in the Equation 3) that must be summed to calculate the total transcription time. This means that there are ten “effective” or “pseudo”-rates. We label these as such because they do not necessarily correspond to actual biochemical reactions but rather could be composed of both the forward and reverse rates, or multiple sub-steps. For example, the effective time the promoter spends trying to move past the first state in the cycle, *T_1_*\* is the sum of four ‘pseudo’ times τ_1_, τ_1_*R_2_*, τ_1_*R_2_R_3_* and τ_1_*R_2_R_3_R_4_*, which is equivalent to saying that there are four irreversible reactions in series that together comprise the transition from state i to state 2 (see example cycle in Figure SiA-C and Supplementary Information).

These equations give an intuition for how the output of a cycle depends its structure. Most importantly, the overall rate of transcription is not simply equivalent to the rate limiting step. Instead, the total transcription time depends on the number and reversibility of all steps in the cycle, and the time spent in any given state depends on the subsequent effective rates. An analogy would be trying to file paperwork in the context of a bureaucratic hierarchy; the number of times you will have the pleasure of meeting civil servant 1 depends on the number of times civil servants 2, 3 and 4 tell you that you’ve filled out your paperwork incorrectly and must try again. This is reflected in the concept of “effective” rates, which account for the structure of the transcription cycle; we thus use effective rates in the formulation of our model below.

### A model based on stochastic chemical kinetics captures both reversible and irreversible steps in transcription

We develop a quantitative model of combinatorial regulation of the transcription cycle based on chemical kinetics. This formalism that can accommodate cycles of arbitrary complexity in terms of connectivity and number of steps; the overall rate of transcription is calculated using a master equation formalism that generates closed-form analytical expressions for the mean and variance of the mRNA distributions (Sanchez & Kondev 2008). Using this formalism one can model the transcription cycle as a series of *N*-1 inactive promoter states leading to state *N*, from which an mRNA molecule is produced as the promoter switches back to state 1, from which the cycle starts again. The promoter transitions stochastically between states *N* and *N*+1 with rate *k_N,N+1_*. We explain this formal model in detail in the Experimental Procedures.

We focus here for illustration purposes on the simple case of combinatorial regulation of a transcription cycle composed of two effective forward rates (*k*_1_ and *k*_2_). Note that this simple case does not require the matrix formalism in order to do the appropriate calculations. However, the insights we derive from this simple case hold true even for more complicated cycles, which can be considered in the matrix formalism above.

In addition to defining the structure of the cycle, we must define how activators affect the component rates, which we define as an *“enhancement”* over the basal rate. In the absence of activators, the reaction proceeds at a “basal” rate, *k_TXN_* (0) = *k_basal_*. To understand the influence of activator A on this basal rate, consider a cycle with a single slow step. In the presence of activator A, this rate increases to *k_TXN_* (A) = *k_basal_* + Δ*k(A)*, where Δ*k(A)* represents the rate *enhancement* (the increase in rate *k*) due to TF A. Similarly, *k_TXN_* (B) = *k_basal_* + Δ*k(B)*. If both A and B are both present and they act independently by accelerating the same single rate in parallel (i.e. each does so through a different, non-interacting mechanism), then the rate *enhancements* will sum together: Δ*k* (A,B) = Δ*k*(*A*) + Δ*k*(*B*). By defining *a* = 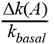 and 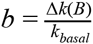, we can express *k_TXN_(A)= k_basal_(1+ a)*, and similarly, *k_TXN_(B)= k_basal_(1+b)*. Therefore, *k_TXN_(A, B)= k_basal_(1+a+b)*, and this defines how in our model, two activators both independently work on the same step.

Now we consider how activators work on the same step, but in the context of a two step cycle. We use *k_1_* and *k_2_* to denote the basal rates of the transitions between state 1→2 and 2→1, respectively, while *a_1_* and *b1* denote the *enhancement* of rate *k_1_* rates by TF A and B, and likewise for rate *k_2_* (*a_i_* ≥ 0 for an activator). Thus, the rate of transcription from a two step cycle in the absence of TFs is

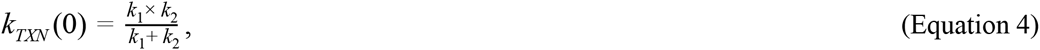

and in the presence of both TFs A and B is

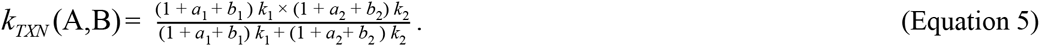

This formulation applies to activators; we use a different formulation for repressors, as described in relation to Figure 2 (see also Experimental Procedures).

**Figure 2.**
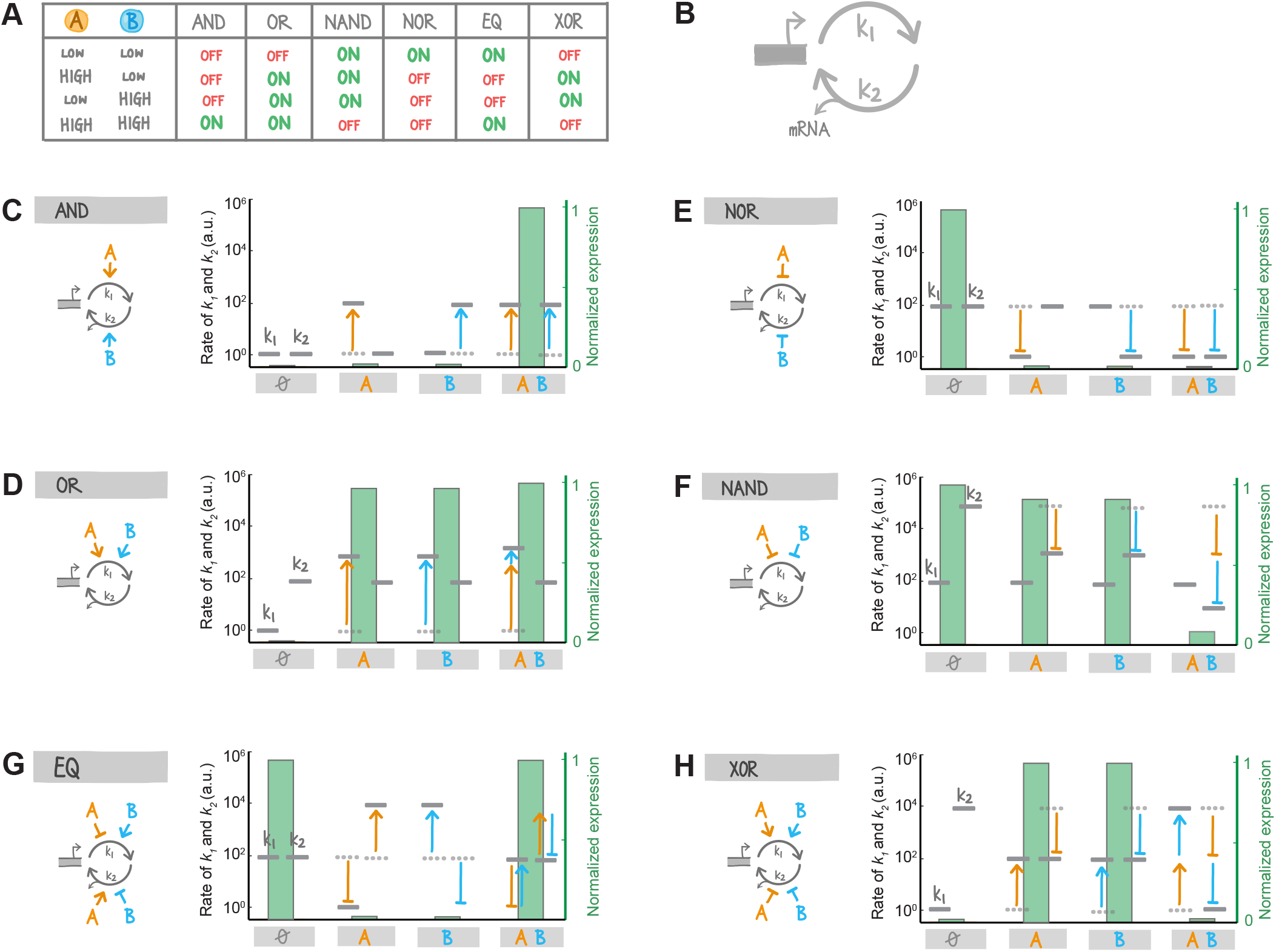
Logical computations could be explained by regulation of the transcription cycle. **(A)** Summary table showing how combinations of TFs A and B map to common Boolean or logical computations. The input TFs are either at low or high concentration, and the output is binary: the gene is ON (highly expressed) or OFF (very lowly expressed, or not expressed at all). **(B)** We consider a simple two-step transcription cycle. The effective forward rates are characterized by the rate constants *k_1_* and *k_2_*, and a molecule of mRNA is produced during step two. **(C-H)** Control of a simple two-step transcription cycle by two TFs can produce all of the basic Boolean computations. For each logic gate we show an example of how TFs A and B could bring about the computation by acting on one or both rates (grey horizontal bars), either as activators (arrows) or repressors (blunted arrows). The resulting expression output due to each TF acting individually or in combination is shown in green, normalised to the maximum expression in each plot such that 0 is “OFF” and 1 is “ON”; equations used to calculate expression in each case are in the Supplementary Information. For the NAND, EQ and XOR gates we assume that when TFs act to reduce the effective rate of their target step, they do so by increasing the free energy of transition; as such, their effects on the rate are multiplicative (see Supplementary Information). The effect of repressor R on a rate is described here as 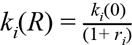, where *r_i_* is the ratio between the repressor concentration and the operator's dissociation constant (*r_i_* ≥ 0).

In the remainder of the results we consider computations that are possible when two TFs act on either or both steps. We assume throughout that TFs work entirely independently of one another; that is, that neither TF helps or hinders the activity of the other. The magnitude of enhancement that an activator has on a given step is determined both by its intrinsic strength and by its concentration. When we randomize the enhancement factor for TFs in our simulations below, this represents randomization of both of these components; thus, we do not explicitly comment on the effect of changing TF concentration alone.

### Logical computations can be achieved by controlling the transcription cycle

Boolean (or logical) computations are commonly used to represent gene regulatory circuits in development and engineering because they produce sharp, on/off decisions (Greber & Fussenegger 2007; Istrail & Davidson 2005; Davidson 2011). Given the importance of these computations at the systems level, it is key to understand how these binary computations may be executed by the cell. Seminal work by Hwa and colleagues addressed this question and found that Boolean logic gates can be implemented for gene regulation by combinatorial recruitment of RNAP (Buchler et al. 2003). These authors demonstrated that all of the elemental two-input logic gates (e.g. AND, OR, NAND, NOR, EQ, XOR) can in principle be built by cooperative pairwise physical interactions between proteins; more complex gates arise by modular arrangement of these pairwise interactions. Here we explore whether logic gates can also be built by regulating the transcription cycle without requiring any physical interactions between TFs, i.e. when activators act independently of one another. Here we use “independent” to mean that the activators neither help nor hinder one another in acting on their shared promoter, in contrast other its alternate usage to describe regulators acting in distinct pathways (i.e. on different promoters).

We calculated the overall rate of transcription by two TFs individually and together, while altering the relative effective rates of the two steps and the strength and direction of TF action (activating or repressing; Figure 2, and Tables S1,2). While there are various ways in which repression could occur mechanistically, here we made the assumption that repressors act by increasing the free energy required to achieve a transition between states; as such, when more than one repressor acts on the same step, their combined effect on that rate is multiplicative. Our analysis shows that the six two-input logic gates described by Buchler et al. (2003) can also be achieved kinetically without invoking any cooperation between TFs. In the simplest example, the AND gate is generated if high expression is possible only when both TFs are present; for this to occur, TFs A and B work as activators each on a different slow step (Figure 2C). In a more complex example, the implementation of the EQ and XOR gates require each TF to activate one step while repressing the other (Figure 2G,H). This dual activation-inhibition effect might for instance occur if, while accelerating one rate, a TF sterically hinders another process in the transcription cycle from taking place.

### Combinatorial control of the transcription cycle allows digital computations by out-of-sync nuclear localization events

Recent experiments demonstrate that TFs can regulate genes by transient bursts of nuclear localization (Cai et al. 2008; Dalal et al. 2014; Hao & O’Shea 2012; Batchelor et al. 2011; Lahav et al. 2004). Furthermore, the timing of colocalization bursts by different regulators may be a means to combinatorially control transcription (Lin et al. 2015). Under the “recruitment” assumption, combinatorially acting TFs must be simultaneously colocalized to the nucleus (i.e. localized in-sync) in order to co-recruit one another and/or the transcriptional machinery (Figure 3A,B) (Lin et al. 2015). In contrast, control of the transcription cycle could enable combinatorial gene regulation even when the bursts of TF nuclear localization are out of sync (Figure 3C, D). All of the logical gates discussed above can be implemented by TFs that are not present in the nucleus at the same time.

**Figure 3.**
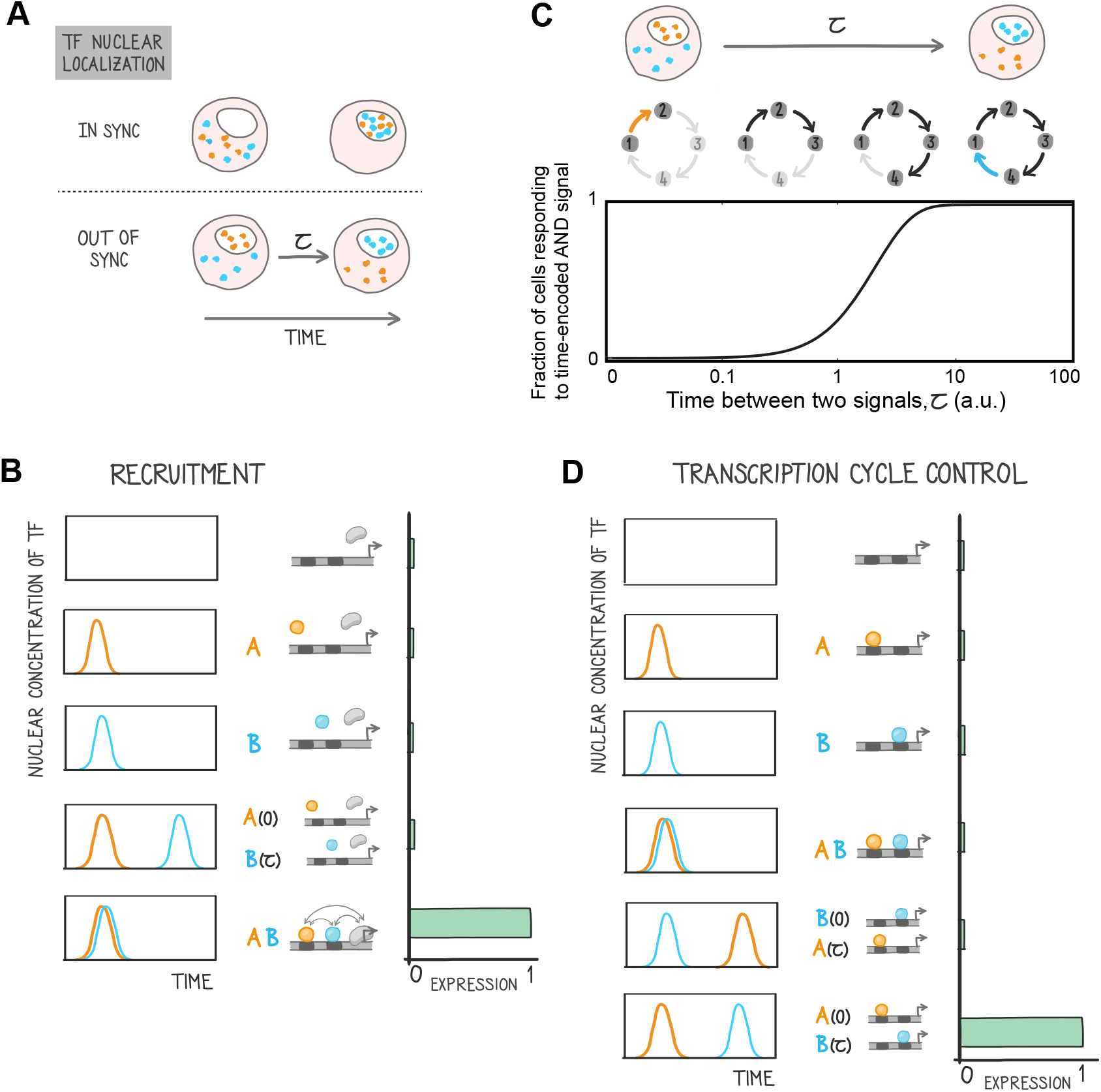
Combinatorial regulation of the transcription cycle enables logical computations on out-of-sync nuclear localization events. **(A)**Recent work has shown that transcription can be combinatorially controlled by transient bursts of transcription factor (TF) localization to the nucleus (Lin et al. 2015). Such localization events could be synchronous (upper panel), or asynchronous (lower panel), with a lag between them of time τ, as shown here for two TFs (orange and blue). **(B)**Under the “recruitment” model of combinatorial transcriptional regulation, TFs must be simultaneously localized to the nucleus in order for a logical computation such as this AND gate to be possible. This is because cooperative physical interactions between the TFs in binding the DNA and/or recruiting RNA polymerase are required to stabilize the binding of RNAP to the promoter. (C, D) In contrast, if transcription is combinatorially regulated by TFs acting on the transcription cycle, the TFs do not need to be on the DNA or even in the nucleus at the same time. We examine a transcription cycle with four effective rates (C, middle panel), which allows the implementation of a time encoded AND gate: An initial localization event of TF A (orange) to the nucleus accelerates the first rate, enabling the subsequent middle two steps in the cycle to occur; these two are slow but proceed without regulation from either TF. After a time delay (τ), TF B localizes to the nucleus and enables completion of the last step in the cycle. With increasing time between the bursts of nuclear localization the fraction of cells responding to this temporally-encoded AND gate increases to 1. **(D)** Temporal encoding of logical computations by regulating distinct steps in the transcription cycle provides a means for the promoter to distinguish the order in which inputs arrive. For this AND-gate example, a high level of expression cannot be achieved if (i) the time required for steps 1, 2 and 3 to occur is longer than the time that TFs A and B are simultaneously colocalized to the nucleus, or if (ii) B arrives before A.

The ability of combinatorial transcription cycle control to work with out-of-sync bursts of TF nuclear localization might provide a means for cells to discriminate the timing of signals. For example, consider two regulators, A and B, that combinatorially activate transcription by acting on a transcription cycle that consists of four slow steps (Figure 3C). TF A accelerates the first step of the cycle (e.g. eviction of a nucleosome) whereas B accelerates the last step (e.g. release of paused RNAP). Two intermediate steps proceed slowly but are unaffected by A or B. TFs A and B are activated by signaling pathways and transiently enter the nucleus in bursts that are separated by time τ. Although the nuclear localization bursts may have different and complex shapes (Cai et al. 2008), for simplicity and analytical tractability we model them as step functions of height *k_A_* and *k_B_* and duration *T_A_* and *T_B_*. We further assume that *k_A_T_A_*, *k_B_T_B_* ≫ 1, such that their respective kinetic step can be completed during nuclear localization of each TF; we also assume the two intermediate steps have equal rates *(k_23_* = *k_34_* = *k_i_)* that are slow compared to the activated rates, such that *k_i_T* ≪1. Qualitatively, none of our conclusions depend on these assumptions, but they simplify the math, allowing us to determine the fraction of cells that respond to this time-encoded AND gate, as a function of the time between the two signals (τ) as:

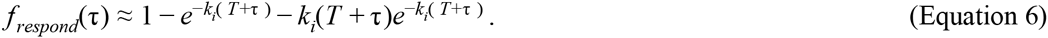

Equation 6 is plotted in Figure 3C. For two temporally-overlapping signals, the cell will not respond to the nuclear localization events Figure (3C,D). Only when the nuclear localization bursts of A and B are shifted by a time larger than τ_*crit*_ = 1/*k_i_* is a significant transcriptional response observed, producing an AND gate. Similarly, if the order of the signals is inverted, and B translocates to the nucleus before A does, the time integrated AND gate produces no transcriptional response (Figure 3D).

This simple example illustrates how kinetic control of the transcription cycle provides a straightforward mechanism by which cells could to add a time component to their logical computations. Combinatorial transcription cycle control could also enable cells to respond to the relative timing between signals (Lin et al. 2015).

### Null model for combinatorial control of transcription: independent action of TFs

Experimental data are rarely entirely digital. Rather, TFs often have analog effects on the level of transcription (e.g. Sharon et al. 2012; Gertz et al. 2009; Patwardhan et al. 2012). To interpret such analog data, we need a null model of combinatorial regulation. Our goal therefore is to calculate the expected behavior of two activators working independently on the same promoter.

Under the recruitment hypothesis, the independent action of TFs on the bound state of RNAP results in multiplicative activation (Veitia 2003; Bintu et al. 2005b; Herschlag & Johnson 1993). This is because each TF additively lowers the energy of the RNAP-bound state of the promoter; since the energies are contained in the exponent of the expression, the effect on mRNA output is multiplicative (Figure 4A and Supplementary Information).

**Figure 4.**
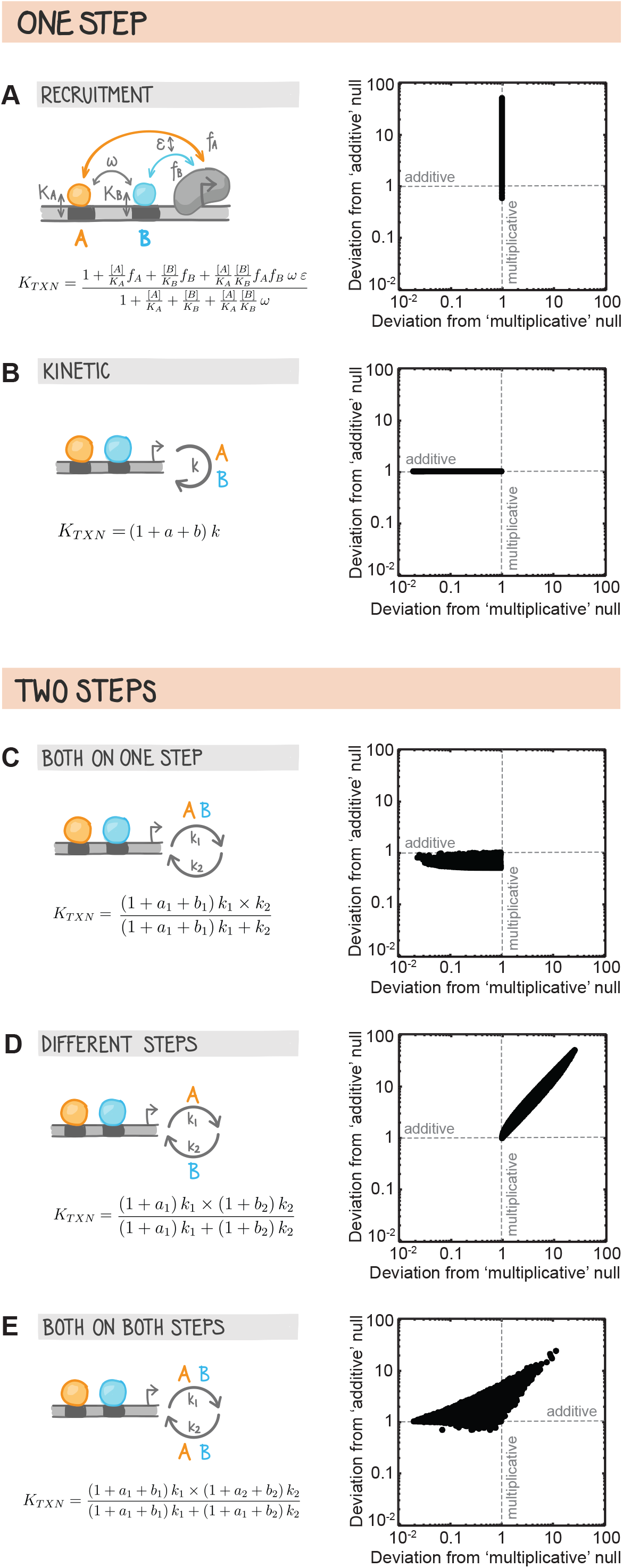
Null models of combinatorial transcriptional control: independent action of activators. To interpret analog data from experiments that examine the effect of transcription factor (TF) number, type and strength on gene expression, we need a null model of combinatorial regulation. Here we define the null model as independent action of TFs. We simulate expression driven by two independently acting activators, using different assumptions about their functions at the promoter. To compare the outcomes, we plot expression relative to the null expectation in two specific limits. In the “recruitment” limit, the null expectation for expression (i.e. expression expected when TFs act independently) is that the *fold changes* in transcription due to each TF are *multiplicative.* On the x-axis we therefore plot deviation from this null, 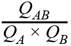, where *Q_TF_* is the fold change in expression over basal. In the “kinetic” limit of two activators working independently on a single slow step, by each working through different pathways, the expectation is that the *enhancement* of the rate by each TF is *additive;* on the y-axis we therefore plot deviation from this null expectation, 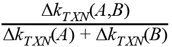. **(A)** Under the “recruitment” hypothesis expression driven by two independently acting activators is expected to be the product of the fold changes due to each TF acting alone. All parameters in the accompanying equation were randomized, except for *ω* and ε, which represent interactions between the activators in binding DNA or contacting the promoter and were held at 1. *K_A_* and *K_B_* are the dissociation constants for TFs A and B; we randomized the parameters 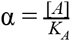 and 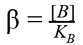 from 0.01-100. *f_A_* and *f_B_* represent the strength of interaction between each TF and the RNA polymerase and these values were randomized from 1-100. **(B)** If either activator can catalyze the rate of the single slow step in transcription via different and independent pathways, the null expectation is that overall enhancement of the rate is the sum of the enhancement due to each TF acting alone. We show this numerically here, where the strength of each activator is randomized between 0-100 while the basal rate, *k,* is varied between 1-1000. (C-E) If there is more than one slow rate in the transcription cycle, the null expectation for expression output is more complex and depends on the function of the activators, i.e. which step(s) in the transcription cycle each acts upon. Simulations show the range of possible gene expression outcomes from a simple cycle composed of two slow steps, *k_1_* and *k*_2_, controlled by two activators, A and B. The strength of enhancement by A and B of each rate is denoted by *a*_1_, *b*_1_, *a*_2_ and *b*_2_, and these were varied between 0-100. This is enhancement strength is determined here both by the intrinsic strength of the activator and by its concentration. *k*_1_ and *k*_2_ were randomized between 1-1000. **(C)** Either activator can catalyze the rate *k*_1_, independently through different pathways. Their enhancement effects on *k*_1_ sum together (see above), but because of the presence of the other rate, *k*_2_, their effects on the change in the overall rate of transcription are largely subadditive, tending to additivity in the limit in which *k_2_ ⋙ k_1_*. **(D)** Two activators working exclusively on different steps in transcription. **(E)** Both activators can work on both steps. For each individual step we assume, as in (B and C), that they catalyze the rate independently through different pathways.

To calculate the effect of independent action of TFs on a transcription cycle, we first consider a simple cycle of only one slow step (Figure 4B). This allows us to compare our results to the recruitment hypothesis, which also implicitly assumes a single rate limiting step. To compare the outcomes of our simulations, we plot the overall rate of transcription in comparison to specific null expectation scenarios (Figure 4). In the “additive” limit, two activators act independently on the same rate limiting step by accelerating it through different pathways; here the expectation is that the transcriptional *enhancements* (Δ*k_TXN_*) caused by each TF add together, so that Δ*k_TXN_(A,B)* = Δ*k_TXN_(A)* + Δ*k_TXN_(B)*. Under the recruitment model, the null expectation is that the *fold-changes* in transcription caused by each TF are multiplicative when both TFs are present. Defining *Q* as the fold-change in transcription over the basal rate due to the presence of an activator, 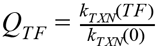, the null expectation for expression in the recruitment limit when both TFs are present is *Q_AB_* = *Q_A_* × *Q_B_*. To visualize how a given mechanism aligns with each null expectation considered above, we construct a null model diagram, where we plot the relative deviation from the “additive” kinetic null model against the relative deviation from the “multiplicative” recruitment null model (Figure 4). Projected on this space, the single-rate recruitment mechanism perfectly aligns with its null, as expected (Figure 4A). The same is true for the single-rate kinetic “additive” mechanism (Figure 4B).

We next consider transcription cycles with two steps, where to formulate a null hypothesis you must choose how TFs will act on the cycle. There are three possible scenarios with two rate limiting steps: TFs accelerate the same step (Figure 4C), different steps (Figure 4D), or both accelerate both steps (Figure 4E). To explore the possible range of outputs one would expect in each case, we randomized the relevant parameters in Equation 5, and projected the results in the null model diagram (Figure 4C-4E). When both TFs work on the same step, the expected distribution is largely sub-additive. The overall transcription rate tends to additivity when the non-regulated step is much faster than the regulated one, such that it has essentially no limiting effect on the rate of transcription. In the second case, when the two TFs work exclusively on different steps, the distribution of rates is always greater than multiplicative. Finally, if both TFs can accelerate both steps, the transcription rate can fall in three of the four quadrants in the null model diagram.

These calculations demonstrate that simple null models of kinetic control over the transcription cycle yield a wide range of possible outputs, depending on the mechanistic assumptions. This emphasizes that, while in some cases additivity emerges as a legitimate null model, this is not necessarily so. Therefore, observation of greater-or less-than-additive expression does not necessarily imply that the underlying mechanisms deviate from independent action of TFs.

Quantitatively formulating null hypotheses is one of the major roles for mathematical modeling in biology. Deviations from the predicted null behavior indicates that there is something surprising happening, either a fault in our understanding of the basic biological mechanisms, or an unidentified interaction between the biological components. In the case of transcription, we have defined our null models as two TFs acting independently from each other. Thus, deviations from the null model would imply non-independent action of TFs, which is our definition of synergy. Synergy can arise through a variety of biophysical mechanisms (Box 2); distinguishing these would therefore be a subject for further study.

##### Box 2: Clarifying the meaning of synergy

The term 'synergy’ arises frequently in discussions of how transcription factors (TFs) work together to affect gene expression. Synergy reflects deviations from a null expectation of independent action of components in a complex system (Salvador 2000). In the context of transcription, because the null expectation is not always rigorously defined, synergy is most often used to mean a “greater-than-additive” effect from multiple bound TFs. Although greater-than-additive effects may arise through various mechanisms described below, synergy itself should not be immediately inferred from it because it depends on the null expectation as we show. There are three main ways that greater-than-additive effects on expression might occur (Herschlag & Johnson 1993; Blau et al. 1996). Notably, two of these reflect null models of independent actions of TFs (and thus cannot actually reflect synergy). This is not an exhaustive list, nor are the mechanisms below mutually exclusive.

**(i) Cooperative DNA binding due to direct or indirect recruitment interaction between TFs.** In this case greater-than-additive effects should disappear at high concentrations of TFs (Carey et al. 1990; Lin et al. 1990).

**(ii) Simultaneous interaction of multiple TFs with the basal transcriptional machinery.** If TFs can simultaneously and independently stabilize the basal transcriptional machinery, their free energies will add, leading to a multiplicative effect on the rate of transcription (Veitia 2003; Herschlag & Johnson 1993). For most values of the individual activator contributions, this will be a greater-than-additive effect (it will not be when the fold-change in transcription for each activator is less than 2).

**(iii) Kinetic control of different steps of the transcription cycle.** If TFs stimulate more than one slow step in transcription then their combined effects will be greater than the sum of their individual actions. This is not the only kinetic means by which greater-than-additive expression could be achieved; for further discussion see Herschlag & Johnson (1993).

**Science Application Box. Predictions arising from combinatorial control of the transcription cycle**

(**A**) Distinguishing functional classes of transcription factors (TFs). Different roles of TFs in stimulating the transcription cycle may be distinguished by assessing the effect of a TF on a promoter that is functionally saturated—i.e. where the expression has plateaued. Adding a binding site for second TF that works on the same step in the cycle (orange) will not increase expression, while one that works on a different slow step will do so if that step is now limiting (red). Calculated expression levels are plotted for a two-state promoter with effective rates *k*_1_ = 1, *k*_2_ = 10 and activator strengths *a_1_*=10, *b_1_* = 2, *c*_2_ =2.

(**B**) The role of the promoter in transcriptional control. Multimerizing binding sites for an activator (orange) that works on one step in a two-step cycle produces different expression depending on the relative rates of the two steps at different promoters (right, promoters A-D). Expression calculated for a two-state promoter with effective rates as shown in the plots to the right, and activator strength *a* = 10.

(**C**) Combinatorial control of transcription by two or more enhancers. Duplicated enhancers may drive little more expression than each enhancer does individually by virtue of responding to the same regulator(s) and thereby accelerating the same rates at the promoter to the same extent. If mutations arise in a duplicate that alter its function, the combined output of the two enhancers may be much higher. This is analogous to some sibling enhancer pairs, which are known to respond to different activators (Wunderlich et al. 2015; Staller et al. 2015). We speculate that these enhancers act differently on the transcription cycle as a result; according to our model, they are thus likely to exhibit super-additive expression. Expression calculated for a transcription cycle with two effective rates, where *k_1_* = *k_2_ =* 1 a.u. and the enhancement factors are *a_1_* = *a’_1_* = 10; *a_2_ = a’_2_* = 2; *a’m_1_* = 2, *a ‘m_2_* = 10.

## BOX: SCIENTIFIC APPLICATION

### Predictions arising from combinatorial control of the transcription cycle

Efforts to uncover a “czs-regulatory code” for transcription—a “grammar” that enables prediction of expression from regulatory sequence—have thus far been only moderately successful. Here we discuss three aspects of transcriptional control that might be illuminated by through a kinetic lens. The kinetic view that we have discussed inspires different types of experiments from the focus on transcription factor (TF) binding site arrangement, and makes concrete predictions as to their outcomes.

#### 1. Distinguishing functional classes of TFs

Little structural information exists for TFs outside the DNA binding domain, making it difficult to assess their function roles. Without structural information, we can consider empirical classification of TFs based on their activity. Recent work elucidated functional classes of TFs by assessing their ability to activate transcription in multiple enhancer contexts (Stampfel et al. 2015). Kinetic control also provides a means of functionally classifying TFs. Multimerizing binding sites for an activator eventually saturates transcription (Sharon et al. 2012; Smith et al. 2013; Burz et al. 1998; Carey et al. 1990). If there is only one slow step in transcription (as assumed by “recruitment” models) this expression level reflects the maximum rate of transcription possible from the promoter in question, and no addition of binding sites for any number of TFs will increase the level of expression above this point. However, if there are multiple slow steps, saturating expression reflects having reached the limiting influence of a different rate. Therefore, while adding a TF that is functionally redundant with the first (i.e. acts on the same step) will have no further effect on transcription, adding one that works on a different slow step will boost transcription (see Figure). Limited experiments of this type have been carried out (Blau et al. 1996), but such experiments also readily adapt to high-throughput screening for functional classes of TFs. It would be valuable to quantitatively assay naturally co-occurring transcriptional activators in order to determine how they control different rates in the transcription cycle.

#### 2. Activator-promoter and enhancer-promoter interactions

Core promoter sequence affects the ability of activators or enhancers to activate transcription (Ohler & Wassarman 2010; Butler & Kadonaga 2001; Zabidi et al. 2015), but this is not incorporated into quantitative models of combinatorial transcriptional control. We anticipate that the spectrum of rates in the transcription cycle differs in a meaningful way between core promoters, according to the existence and arrangement of core elements (such as the TATA box, DPE, and Inr). In more extreme cases a promoter could determine whether an activator is able to act on a rate at all, i.e. whether the two are ‘biochemically compatible’ (Juven-Gershon et al. 2008; van Arensbergen et al. 2014). The model we present provides a quantitative framework in which to interpret differences in the expression from enhancer-promoter or activator-promoter pairs. We illustrate this in the accompanying Figure by plotting the expression levels driven by multimerized activator binding sites at four hypothetical core promoters. Because the promoters differ in the relative rates of the two slow steps in this transcription cycle, the effect that any given number of bound activator molecules has on the level of gene expression differs between them. Similarly, the effect of an enhancer bound by activators that work only on the slowest step in transcription would vary widely depending on the ratio between this and next-slowest step. By pairing enhancers with libraries of promoters that have been classified by their core element content (Ohler et al. 2002) it may be possible to work backwards to deduce enhancers’ roles in stimulating the transcription cycle.

**Box: Scientific Application.**
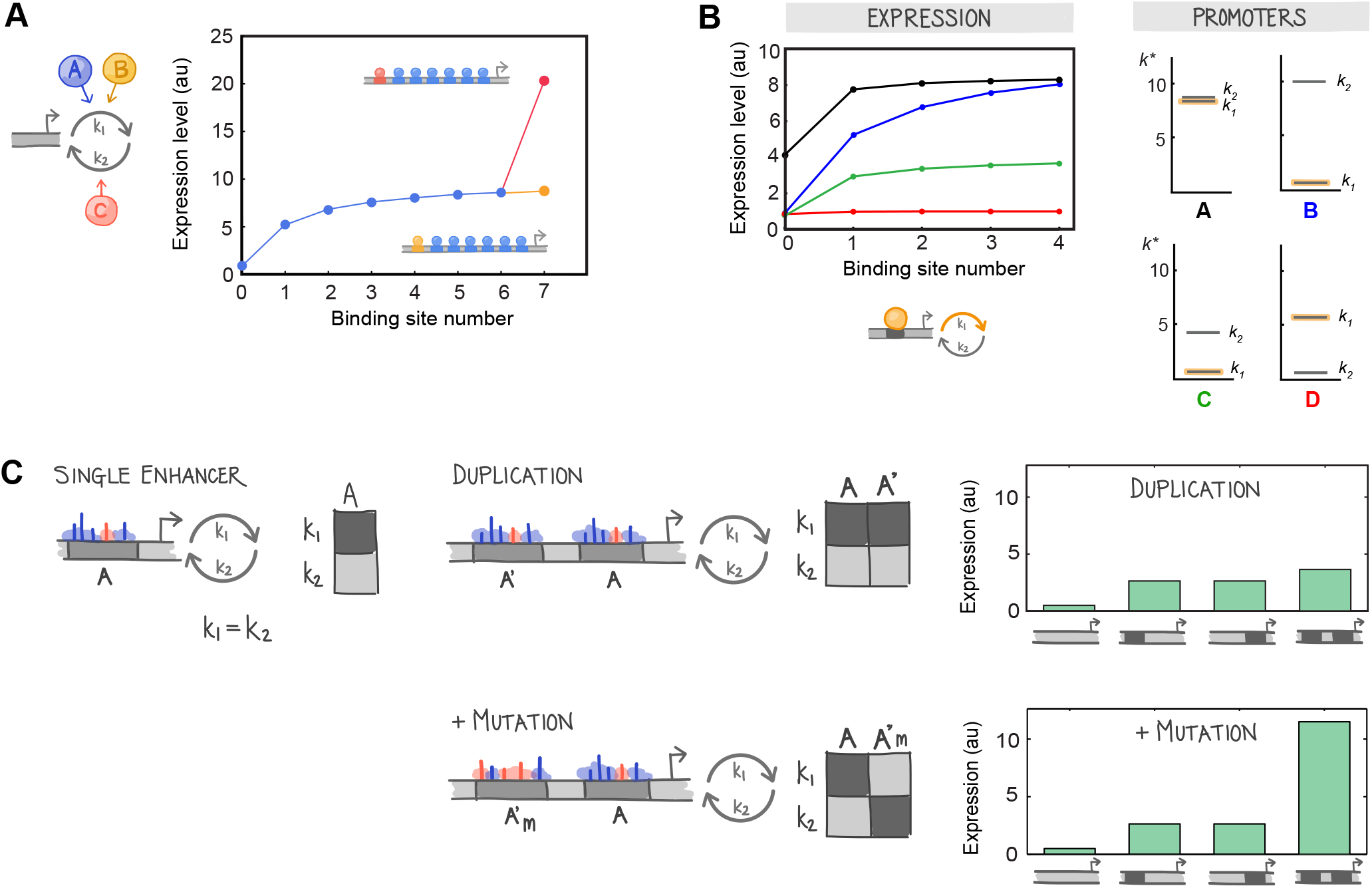
Predictions arising from combinatorial control of the transcription cycle. **(A)** Distinguishing functional classes of transcription factors (TFs). Different roles of TFs in stimulating the transcription cycle may be distinguished by assessing the effect of a TF on a promoter that is functionally saturated—i.e. where the expression has plateaued. Adding a binding site for second TF that works on the same step in the cycle (orange) will not increase expression, while one that works on a different slow step will do so if that step is now limiting (red). Calculated expression levels are plotted for a two-state promoter, with effective rates *k_1_=1, k_2_=10* and activator strengths *a_1_* = 10, *b_1_=* 2, c_2_= 2. **(B)** The role of the promoter in transcriptional control. Multimerizing binding sites for an activator (orange) that works on one step in a two-step cycle produces different expression depending on the relative rates of the two steps at different promoters (right, promoters A-D). Expression calculated for a two-state promoter with effective rates as shown in the plots to the right, and activator strength a = 10. **(C)** Combinatorial control of transcription by two or more enhancers. Duplicated enhancers may drive little more expression than each enhancer does individually by virtue of responding to the same regulator(s) and thereby accelerating the same rates at the promoter to the same extent. If mutations arise in a duplicate that alter its function, the combined output of the two enhancers may be much higher. Expression calculated for a transcription cycle with two effective rates, where *k_1_* = *k_2_* = 1 a.u. and the enhancement factors are *a_1_* = *a’_1_* = 10; *a_2_* = *a’_2_* = 2; *a’m_1_* = 2, *a’m_2_* = 10.

#### 3. Combinatorial control by two or more enhancers

Many developmental genes contain two or more enhancers that drive spatially and temporally overlapping patterns of expression from the same promoter (Frankel et al. 2010; Hoch et al. 1990; Dunipace et al. 2011; Wunderlich et al. 2015; Staller et al. 2015; Barolo 2011; Prazak et al. 2010; Perry et al. 2011). The mechanism by which the information from these “shadow” enhancers is combined in single cells to drive a given rate of transcription is unknown and the combined output is unpredictable (Dunipace et al. 2011; Bothma et al. 2015; Mas et al. 2004). It is plausible that shadow enhancers work together by regulating different steps in the transcription cycle. The strength of effect that an enhancer has on each rate in the transcription cycle will depend on the number and type of TFs bound to it in any given cell. But since TF concentrations change over space and time, so too may the the enhancer's effect on the transcription cycle and how the expression it drives is integrated with that from other enhancers. The common experiment to test for interactions between enhancers (measuring the influence of each alone and together) is difficult to rigorously interpret since multiple models could explain the experimental data (see Figure 4). A variation on this experiment that may prove useful is to compare the effects of enhancer duplications to those of shadow enhancers. We speculate that a pair of shadow enhancers may behave differently than a duplication of either. Duplicate enhancers contain an identical number and type of transcriptional activators, which makes them likely to work in the same manner on the transcription cycle, leading to a combined expression output that is largely subadditive. If mutations accumulate in a duplicated pair of enhancers their functions could diverge, leading to different effects on transcription. This is analogous to some sibling enhancer pairs, which are known to respond to different activators (Wunderlich et al. 2015; Staller et al. 2015).

## DISCUSSION

We argue that a quantitative model of combinatorial gene regulation should include multiple steps in the transcription cycle. We show that two TFs acting independently can produce a wide range of outputs, both analog and digital. These outputs depend on the underlying rates of the transcription cycle and how the TFs act on these rates. This view of transcriptional control emphasizes the information-processing role of the promoter, which dictates the spectrum of effective rates in the transcription cycle. We speculate that combinatorial control of the transcription cycle enables greater plasticity than a process that relies only on physical interactions between TFs; this could enable rapid regulatory evolution as is observed in many systems.

### A cis-regulatory “code” at the level of TF function?

The holy grail of deciphering regulatory DNA is unlocking the “cis-regulatory code”, which ideally would allow us to predict gene expression driven by any regulatory sequence. The emphasis in efforts to crack this code has been on the number, affinity and arrangement of TF binding sites in enhancers (reviewed in Weingarten-Gabbay & Segal 2014). This emphasis is reasonable if combinatorial control of transcription is channeled through regulation of a single slow step in the transcription cycle. However, while binding site arrangement does play an important role in cooperative DNA binding (Thanos & Maniatis 1995; Adams & Workman 1995), a general code at the level of TF binding sites has remained elusive (Weingarten-Gabbay & Segal 2014). We have shown here that kinetic control of the transcription cycle alone can also produce complex computations. From this kinetic perspective, it may matter less how TF binding sites are configured and matter more which TFs are recruited to regulatory DNA. Recent work categorizes TFs into functional classes based on their combinatorial activity in different enhancer contexts (Stampfel et al. 2015), which offers the opportunity to test the kinetic perspective directly.

Kinetic control of the transcription cycle casts the basal promoter in a key role. While some communities recognize the information processing role of the promoter (Ohler & Wassarman 2010; Danino et al. 2015; Zehavi et al. 2014), others consider enhancers the source of spatiotemporal patterns of expression, while basal promoters are a passive elements that make genes competent for transcription to a greater or lesser extent (“strong” versus “weak” promoters). We show that TF function is necessarily interpreted through the rate-limiting steps of the promoter, firmly putting the promoter in an information processing role in a quantitative modeling framework. This framework can now accommodate multiple experimental observations. For example, the composition of core motifs in basal promoters plays a key role in cell type-specific gene regulation (reviewed in (Danino et al. 2015; Zehavi et al. 2014; Ohler & Wassarman 2010)), affecting the composition of the preinitiation complex (Lewis et al. 2005), pausing (Amir-Zilberstein et al. 2007; Hendrix et al. 2008), and termination and recycling of RNAP (Mapendano et al. 2010). Moreover, activation domains have different abilities to work with different core promoter elements, emphasizing how these two types of regulatory DNA work together (Emami et al. 1995; Das et al. 1995; Juven-Gershon et al. 2008).

### The model of transcription cycle control readily accommodates endogenous complexities

Building models necessarily involves deciding which features to include and which to leave out. Our goal in this paper has been to highlight how combinatorial regulation of the transcription cycle may be exploited by cells to process information, an area that has received relatively little attention. To accomplish this we purposely simplified the mechanisms by which TFs bind, unbind and interact with the transcriptional machinery. We used a mean field approximation, where the effect of TFs on individual rates were described by their “activity”, which corresponds to the product of the rate of activation when a TF is bound and the probability that the TF is indeed bound. The latter is a function of its concentration, binding affinity and mode of binding (independent, cooperative, etc.). In practice, activation by a TF is a single molecule stochastic process and thus our approach is only an approximation that may very well not capture the actual biophysics in many instances. However, while these details are important on a case-by-case basis, their abstraction into a single effective parameter is what allowed us to examine the transcription cycle in isolation. We note that a full model incorporating both the stochastic TF association/dissociation and the mechanistic step of accelerating the rates that they regulate can be easily accomplished using the same formalism that we employed here.

For the purposes of illustration we considered a very simple scenario of combinatorial control of the transcription cycle consisting of one or two slow steps. Of course, transcription is much more complex. For example, promoters differ in which and how many rates are limiting. The model we describe is agnostic to the identities of the slow steps in a transcription cycle. What determines the overall rate of transcription is the relative rates and how TFs impinge upon them, both of which can be computationally explored. There is also not a one-to-one mapping of TF to function, i.e. a single TF can work on multiple steps either directly or via the cofactors with which it interacts (reviewed in (Fuda et al. 2009)). For example, the yeast activator Gcn4 contacts four coregulator complexes, each of which affect chromatin remodeling, assembly of the preinitiation complex and elongation to different degrees (Fishburn et al. 2005; Govind et al. 2005). Reciprocally, a single process can be influenced by multiple TFs, often through shared intermediates like Mediator. In our model, the catalysis of a given rate can be arbitrarily apportioned between TFs, and vice versa. Finally, the concentrations and cooperativities in binding to DNA will also play roles in determining the outcome of transcription. Equilibrium models based on statistical thermodynamics have already explored these in detail (Bintu, Buchler, Garcia, Gerland, Hwa, Kondev, Kuhlman, et al. 2005; Bintu, Buchler, Garcia, Gerland, Hwa, Kondev & Phillips 2005). The purpose of our analysis has been to complement these models with the kinetic aspects that are most apparent when the binding sites are already saturated.

### Control of the transcription cycle may enable evolutionary plasticity

Here we have addressed how a promoter could execute computations without relying on interactions between transcription factors. But an interesting question going forwards will be what the advantages and disadvantages are of generating computations through TF interactions, versus by controlling different kinetic steps in transcription, with respect to how regulatory evolution can occur. We speculate that control of the transcription cycle could help to enable the flexibility apparent in how gene expression patterns are encoded in regulatory DNA between species (Ludwig et al. 2005; Wunderlich et al. 2015; Tuch et al. 2008; Tsong et al. 2006). As we have discussed, by combinatorially controlling the transcription cycle, the the same computations can be encoded without the need for any cooperation between transcription factors. This relieves the need for interacting TF binding sites to be in close vicinity with one another, removing a significant constraint on how binding sites are organised within regulatory sequences. In addition, we suggest that kinetic control might also enable changes to accumulate in regulatory DNA through “buffering” their potential phenotypic consequences. Binding sites for TFs that work on fast rates may come and go without affecting gene regulation, but such variation could be unmasked by mutations in promoter sequence, or environmental or physiological changes. This is analogous to the evolutionary capacitor function of molecular chaperones (Rutherford & Lindquist 1998).

### Outlook

Given the overwhelming evidence for action of TFs on multiple distinct, slow steps in transcription, it is time to incorporate this reality into models of combinatorial gene regulation. Viewing transcriptional regulation through a kinetic lens may help to unify and explain multiple experimental realities, such as complex transcription cycles, fast TF binding, and flexible regulatory evolution. In the long term, elucidating TF function beyond simple ‘activation’ or ‘repression’ may finally reveal a tractable *cis*-regulatory code.

## EXPERIMENTAL PROCEDURES

### Calculating mean mRNA production from an arbitrarily complex promoter

Due to the stochastic nature of transcription, the instantaneous rate of initiation will fluctuate over time, even under a constant environment. To calculate the moments of the distributions of transcription rates within a single cell over time (which, by assuming ergodicity also reflect the moments of the distribution of initiation rates within a population of genetically homogeneous cells exposed to the same environment), we use a matrix formalism akin to that which we introduced in previous work (Sanchez & Kondev 2008; Sanchez et al. 2011; Sanchez et al. 2013), inspired by earlier models of a two-state promoter (Peccoud & Ycart 1995; Kepler & Elston 2001).

To illustrate this formalism, consider a promoter that can exist in three states. Transitions rates from state *i* to *j* is described by *k_ij_*. We assume in this example that the only transitions allowed are: from 1 and 2 (in both directions, described by *k*_12_ and *k*_21_), from 2 and 3 (in both directions: *k*_23_ and *k*_32_), and finally that a productive transition (i.e. one that is accompanied by the synthesis of a new mRNA molecule) is allowed from 3 to 1 (given by *k*_31_); this last transition is irreversible. We introduce a vector containing the steady state probabilities of finding the promoter in states 1, 2 and 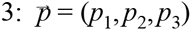. The average rate of transcription will be equal to the dot product of the vector of mRNA synthesis from each state 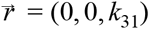 and the probability vector 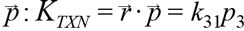. In order to calculate how *p* depends on the kinetic rates, we just need to solve the equation: 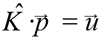, where 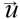 is the vector of the same dimension as 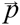 (which we’ll define as *L*), and whose entries are *u_j_* = δ*_jL_*. In turn, 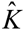 is the matrix with entries 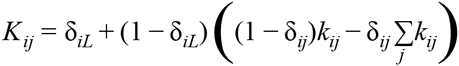. In both of these equations *δ_ij_* represents the Kronecker delta, i.e. δ*_ij_* = 1 if *i=j,* and δ*_ij_* = 0 otherwise.

### Combinatorial regulation of transcription by bursts of nuclear localization

The fraction of cells that have produced a transcript in response to the nuclear localization of A and B for a four step kinetic cycle such as that depicted in Figure 3 can be determined as follows. First, we define the promoter state vector 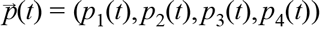, which contains the probabilities that a promoter will be at each state (or what is the same, the fraction of cells in a population that are at those states) at time *t*. To calculate the fraction of cells that have produced at least one transcript in response to the nuclear localization events, we consider the transition from state 4 to state 1 (which produces a transcript) as leading to an absorbing boundary (this is a convenient mathematical artifice with no bearing on the actual biochemistry). The value of the vector 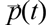 can be found by solving the equation: 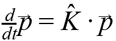, where 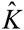 in this case has elements: *K*_11_ = −*K*_21_ = − *k*_12_, *K*_22_ = − *K*_32_ = − *k*_23_, *K*_33_ = − *K*_43_ = − *k*_34_and *K*_44_ = − *K*_14_ = − *k*_41_. For all other values of *i,j*, *K_ij_* = 0. In the example we worked out in the main text, we assumed that the first step in the cycle, *k*_12_, depends on the intranuclear concentration of A, whereas the final step in the cycle, *k*_14_, depends on the intranuclear concentration of B. Since both [*A*] and [*B*] change with time as a result of a nuclear localization burst, this leads to time dependent functions which we write as: *k*_12_ = *f_A_(t)*, *k*_41_ = *f_B_(t)*. In order to investigate an example of this model that can be solved analytically, in simple terms but without losing generality, we assume that *f_A_(t)* is such that it causes essentially all cells to complete the first step of the cycle in a very short time (in comparison to the time required to complete the second step of the cycle). In practice, this can be modeled by a square step function with very large height (*k_A_*) and very narrow width (*T*), so that *k_A_T ≫ 1 ≫ k_23_T*. Likewise, we also assume that *f_B_(t)* has the same shape as *f_A_(t)*, and we also assume the transitions from state 2 to state 3, and from state 3 to state 4 both happen with the same rate *k_i_*, which is independent of the intranuclear concentrations of A and B. Solving this system of equations under the constraints and assumptions discussed above yields Equation 6 in the text

### Simulating analog and digital computations through control of the transcription cycle

To explore the space of mean transcription rates that could be achieved under a given model of activator function on a simple two-step cycle (Figure 4), we randomized key parameters in Equation 5 in the main text. Specifically, we randomized the rates of the two slow steps in the cycle over three orders of magnitude and varied activator strength, described by {*a_1_, a_2_, b_1_, b_2_*}, between 0 and 100; this is within the same order of magnitude as the measured effect of an activator on a given rate in transcription (Friedman & Gelles 2012). To explore the transcription rates that can be produced under the recruitment model, we randomized the parameters in the equation shown in Figure 4A (Bintu et al. 2005a). Because we were interested only in the null situation of activator independence, we held ω and ε at 1 (these describe cooperativity between TFs in binding to DNA and in contacting RNAP, respectively). The concentrations of activators A and B ([*A*] and [*B*]) were varied between 0.01 - 100, while *f_A_* and *f_B_* were varied between 1 -100.

To investigate potential means of achieving logical calculations from a cycle of two slow effective rates controlled by two transcription factors we surveyed a range of parameters and chose a set that worked well for illustration. The parameter values used are listed in Table S1.

## AUTHOR CONTRIBUTIONS

C.S., A.H.D and A.S. conceived and designed the study and wrote the manuscript. C.S. and A.S. developed the analytical framework.

## ACKNOWLEDGEMENTS

We would like to thank Hernan Garcia, Jané Kondev and Sandeep Choubey for many useful discussions, and Stirling Churchman, Jeremy Gunawardena, Mike Springer, Johan Paulsson and Jané Kondev for their comments and feedback on the manuscript. A.S. was supported by a Rowland Junior Fellowship from The Rowland Institute at Harvard. C.S. was supported by a Harvard Graduate Society Research Fellowship. Work in the DePace lab is supported by NIH U01 GM103804-01A1, NSF CAREER IOS-1452557, and the Grinnell Fund for Biomedical Research.

## SUPPLEMENTARY INFORMATION

### Calculating effective rates from a closed-loop cycle with reversible and irreversible steps

Refers to Box 1 and first section of Results, and accompanies Figure S1.

In Box 1 we showed two examples of the 'spectra’ of effective rates at a bacterial and a eukaryotic promoter. We calculated these effective rates, which incorporate the effects of reverse rates, using kinetic data from (Friedman & Gelles 2012)) and (Stasevich et al. 2014)). Here we describe how we obtained these effective rates, which we refer to as ‘effective’ because they do not necessarily correspond to actual biochemical reactions.

The transcription cycle at the activator-dependent σ^54^ promoter of the *S. typhimurium* glnALG operon is comprised of reversible and irreversible transitions between four principal states (Friedman & Gelles 2012) (Figure S1A). State 1 is the unbound promoter, states 2 and 3 are closed complexes, and state 4 is the open complex; mRNA is produced from the elongation step (rate *k_41_*). The forward and reverse rates connecting the four states were measured using multiwavelength single-molecule fluorescence colocalization (CoSMoS) (Friedman & Gelles 2012). The rate of RNAP association with the promoter, *k_12_*, was determined to be 2.1 × 10^7^ M^-1^ s^-1^; we assumed that RNAP exists at ∽1uM in the cell, giving a rate of rate of 21 s^-1^, which we use in these calculations. In the absence of the bacterial activator, *k_34_* is < 6 × 10^-6^; this rate increases 300-fold to 1.93 × 10^-3^ s^-1^ when the activator NtrC is added along with ATP, but the rates spectrum shown here is in the absence of the activator.

This four-step cycle (Figure S1B) is comprised of ten constituent ‘pseudo’-times that are summed together to calculate the total time taken to complete the cycle *(T_TXN_*; Equation 3, main text). These terms do not correspond to real biochemical reactions. For example, the effective time the promoter spends trying to move past the first state in the cycle, *T_t_*\* is the sum of four ‘pseudo’ times *τ_1_, τ_1_R_2_, τ_1_R_2_R_3_* and τ_1_*R_2_R_3_R_4_*, where R is the rejection rate (Equation 1, main text), which is equivalent to saying that there are four irreversible reactions in series. The ten pseudo-steps are diagrammed in Figure S1B and their rates (the inverse of the time taken to complete each pseudo-step) are plotted in Figure S1C, color-coded by the state transition to which they belong.

For the eukaryotic transcription cycle example we used kinetic measurements of rates in the transcription cycle made by Kimura and colleagues on a glucocorticoid receptor activated tandem gene array in a mouse cell line. These authors developed a means of tracking specific histone modifications and RNAP phosphorylation in living cells and used these to measure the association of RNAP with the promoter in the presence of GR (state 1 → state 2, Figure S1D), promoter escape (state 2 → state 3; associated with phosphorylation of serine 5 of the RNAP C-terminal domain), and escape into sustained elongation (state 3 → state 1; associated with ser2 phosphorylation of the C-terminal domain). The corresponding forward and reverse rates are detailed in Figure S1D.

**Figure S1.**
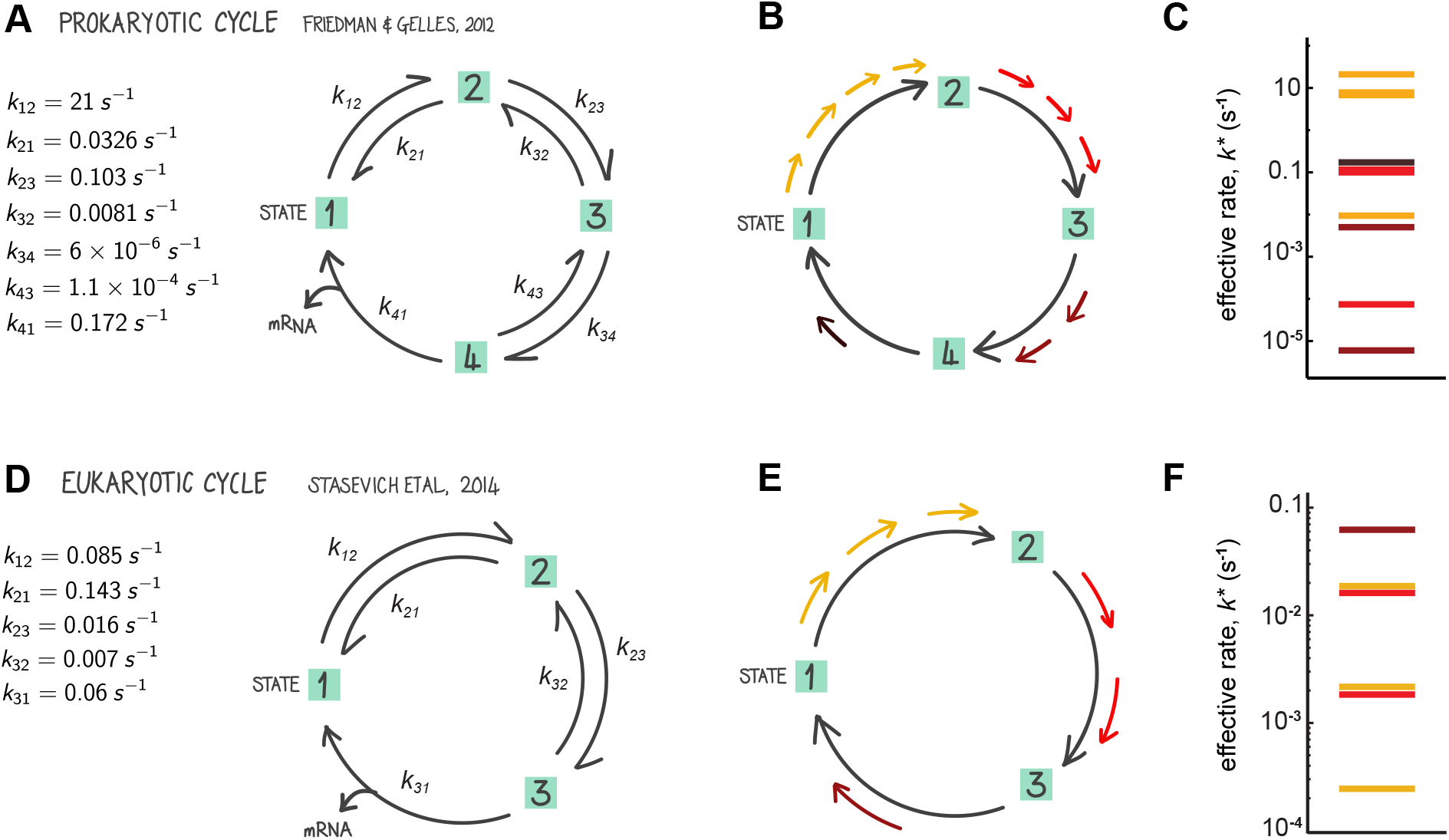
Calculating effective rates from a closed-loop cycle with reversible and irreversible steps. This Figure accompanies the rates “spectra” in Box 1 and the explanation of calculating the transcription rate from a cycle with no internal loops in the Results section. The forward and reverse rates for a prokaryotic and a eukaryotic transcription cycle were obtained from Friedman & Gelles (2012) and Stasevich et al (2014), respectively. These are listed next to a simplified representation of each cycle in (A) and (D). As described in the accompanying text, we calculated the effective forward rates for each cycle; these take into account the effect of reverse rates. For example, in (A-C) the promoter transitions between four states; the total time taken to complete the cycle, *T_TOTAL_*, is the sum of the times taken to complete each step. For the four-step cycle in question, *T_TOTAL_* comprises ten terms (see Results), and the inverse of each term corresponds to an ‘effective’ rate. We refer to them as ‘effective’ because they don’t necessarily correspond to a real biochemical reaction since they incorporate the effects of all the reverse rates in the cycle). In (C) and (F) we plot the effective rates (*k*\*) for each cycl

**Assumptions made in examples of logic gate formation through control of the transcription cycle**

**Table S1.** Parameter values used to generate logic gates in Figure 2.

|  | AND | OR | NOR | NAND | EQ | XOR |
| --- | --- | --- | --- | --- | --- | --- |
| $k_1$ | 1 | 1 | 100 | 100 | 100 | 1 |
| $k_2$ | 1 | 100 | 100 | 100 000 | 100 | 10 000 |
| $a_1$ | 100 | 1000 | 0.01 | 0 | 100 | 100 |
| $b_1$ | 0 | 1000 | 0 | 0 | 0.01 | 100 |
| $a_2$ | 0 | 0 | 0 | 0.01 | 0.01 | 0.01 |
| $b_2$ | 100 | 0 | 0.01 | 0.01 | 100 | 0.01 |

**Supplementary Table 2:** Calculating rates of transcription in Figure 2. Equations used to calculate the rate of transcription for the example logic gates shown in Figure 2. We consider a simple cycle comprised of two steps, described by rate constants, *k_1_* and *k_2_*. *k_TXN_*(0), overall rate of transcription in the absence of activators; *k_TXN_*(A), transcription rate in the presence of activator A; *k_TXN_*(AB), transcription rate when both activators A and B are present. *a_1_* and *a_2_* represent the enhancement of rates *k_1_* and k_2_, respectively, by activator A. When an activator has no effect on a rate we have left it out of the equations for clarity.

|  | AND | OR | NOR |
| --- | --- | --- | --- |
| $k_{TXN}(0)$ | $\frac{k_1 \times k_2}{k_1 + k_2}$ | $\frac{k_1 \times k_2}{k_1 + k_2}$ | $\frac{k_1 \times k_2}{k_1 + k_2}$ |
| $k_{TXN}(A)$ | $\frac{(a_1 + 1) k_1 \times k_2}{(a_1 + 1) k_1 + k_2}$ | $\frac{(a_1 + 1) k_1 \times k_2}{(a_1 + 1) k_1 + k_2}$ | $\frac{(a_1) k_1 \times k_2}{(a_1) k_1 + k_2}$ |
| $k_{TXN}(B)$ | $\frac{k_1 \times (b_2 + 1) k_2}{k_1 + (b_2 + 1) k_2}$ | $\frac{(b_1 + 1) k_1 \times k_2}{(b_1 + 1) k_1 + k_2}$ | $\frac{k_1 \times (b_2) k_2}{k_1 + (b_2) k_2}$ |
| $k_{TXN}(A,B)$ | $\frac{(a_1 + 1) k_1 \times (b_2 + 1) k_2}{(a_1 + 1) k_1 + (b_2 + 1) k_2}$ | $\frac{(a_1 + b_1 + 1) k_1 \times k_2}{(a_1 + b_1 + 1) k_1 + k_2}$ | $\frac{(a_1) k_1 \times (b_2) k_2}{(a_1) k_1 + (b_2) k_2}$ |

|  | NAND | EQ | XOR |
| --- | --- | --- | --- |
| $k_{TXN}(0)$ | $\frac{k_1 \times k_2}{k_1 + k_2}$ | $\frac{k_1 \times k_2}{k_1 + k_2}$ | $\frac{k_1 \times k_2}{k_1 + k_2}$ |
| $k_{TXN}(A)$ | $\frac{k_1 \times (a_2) k_2}{k_1 + (a_2) k_2}$ | $\frac{(a_1 + 1) k_1 \times (a_2) k_2}{(a_1 + 1) k_1 + (a_2) k_2}$ | $\frac{(a_1 + 1) k_1 \times (a_2) k_2}{(a_1 + 1) k_1 + (a_2) k_2}$ |
| $k_{TXN}(B)$ | $\frac{k_1 \times (b_2) k_2}{k_1 + (b_2) k_2}$ | $\frac{(b_1) k_1 \times (b_2 + 1) k_2}{(b_1) k_1 + (b_2 + 1) k_2}$ | $\frac{(b_1 + 1) k_1 \times (b_2) k_2}{(b_1 + 1) k_1 + (b_2) k_2}$ |
| $k_{TXN}(A,B)$ | $\frac{k_1 \times (a_2 \times b_2) k_2}{k_1 + (a_2 \times b_2) k_2}$ | $\frac{b_1(a_1 + 1) k_1 \times a_2(b_2 + 1) k_2}{b_1(a_1 + 1) k_1 + a_2(b_2 + 1) k_2}$ | $\frac{(a_1 \times b_1 + 1) k_1 \times (a_2 \times b_2) k_2}{(a_1 \times b_1 + 1) k_1 + (a_2 \times b_2) k_2}$ |

There are many potential ways in which the logic gates in Figure 3 might be achieved mechanistically. We focus only on examples of necessary basal rates in the transcription cycle and changes from them. In the scenarios in Figure 3 we assumed the following. AND gate: Each TF independently catalyzes a different rate. OR gate: Either TF can independently catalyze the slower rate, therefore their effects on this rate add together. NOR gate: Each TF acts as a repressor on a different rate. The effect of repressor R on a rate is described here as 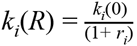, where *r_i_* is the ratio between the repressor concentration and the operator's dissociation constant ( *r_i_* ≥ 0). NAND gate: Both TFs can independently repress the faster of the two rates. We assume for the NAND and EQ gates that repressors act by increasing the free energy required to complete a step. Therefore, when they work independently on the same step, their combined effects on this rate of this step are multiplicative. EQ gate: We assume that each TF activates one rate and represses the other. This might for example occur if, while carrying out its activating function at rate 1, the TF occludes recruitment of a key enzyme necessary for step 2 to proceed. XOR gate: Both TFs increase the slower rate while decreasing the faster rate. When both are present, their combined effects on each rate are multiplicative. This could occur if the TFs act by stabilizing or destabilizing a given protein complex.

### Under the recruitment model, expression is multiplicative when TFs act independently

The null expectation for expression controlled by two activators acting independently under the recruitment model is multiplicativity (Bintu et al. 2005b; Veitia 2003). This has been derived analytically, which we recap below to accompany Figure 4A, but we also demonstrate it numerically in that Figure.

In the recruitment model, activators recruit RNAP to the promoter by lowering the free energy of the bound RNAP state. The rate limiting step is the recruitment of RNAP because *k_OFF_*is extremely fast, such that RNAP falls off the promoter rapidly before getting the chance to fire. Activators work by reducing this off-rate. We assume that activators A and B have affinities described by the dissociation constants *K_A_* = *e*^−Δ*G*_*A*_^ and *K_B_* = *e^−Δ*G*_B_^*, where Δ*G_A_* and Δ*G_B_* represent the binding free energies for the two activators. We consider activators A and B to be present at concentrations [*A*] and [*B*]. When bound at the same time, both activators stabilize each other by reducing their dissociation constants by a factor ω = *e*^−Δ*G_AB_*^. A and B also lower the free energy of the bound RNAP state by Δ*G_AP_* and Δ*G_BP_* respectively, and they thus decrease the dissociation constant of RNAP by magnitudes *f_A_* = *e^−ΔG_Ap_^* and *f_B_* = *e^−ΔG_Bp_^*. If the two TFs contact the RNAP independently from each other, the net reduction in free energy of the bound RNAP will be the sum of the reductions caused by each TF. If they do not act independently, the extent of the deviation is described by a parameter ε = *e^−g_AB_^*, so that 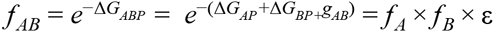. When only the binding site of A is present (and defining for convenience the unitless parameter 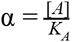), the fold-change activation takes the form 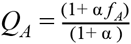. Likewise, and defining 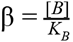, the fold-change activation when only the binding site of B is present is 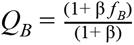. The fold-change activation of when both binding sites are present takes the form 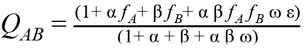.

These results can be easily derived from the model, but they can be also found elsewhere (Bintu, Buchler, Garcia, Gerland, Hwa, Kondev, Kuhlman, et al. 2005; Bintu, Buchler, Garcia, Gerland, Hwa, Kondev & Phillips 2005). In Figure 4A, since we are interested in the null situation of independent action of activators (i.e. *g_AB_=* 0, Δ*G_AB_=* 0), we set ε =1, ω = 1 in the model, and randomized all of the other parameters in the above equation for *Q_A_, Q_B_*, and *Q_AB_.*

### Under the recruitment model, additivity is rare

Under the recruitment model, in order to get additive effects between two binding sites when the activators act independently to recruit RNAP, it is necessary that all of the following conditions are fulfilled (refer to the equation in the previous section): (i) α << 1, (ii)β << 1, (iii)*f_A_x* α >> 1, (iv)*f_B_x* β >> 1 and (v) either ω = 0 or ε = 0; i.e. the activators bind to their binding sites only rarely but have very strong effects on the RNAP when they are bound, and they cannot simultaneously contact RNAP. Therefore, while additivity is in principle possible under the recruitment mechanism, it can only occur under very restricted conditions; therefore the recruitment mechanism is unlikely to yield additive behavior.

